# Formulation excipients and their role in insulin stability and association state in formulation

**DOI:** 10.1101/2022.08.01.502380

**Authors:** Caitlin L. Maikawa, Leslee T. Nguyen, Joseph L. Mann, Eric A. Appel

## Abstract

While excipients are often overlooked as the “inactive” ingredients in pharmaceutical formulations, they often play a critical role in protein stability and absorption kinetics. Recent work has identified an ultrafast absorbing insulin formulation that is the result of excipient modifications. Specifically, the insulin monomer can be isolated by replacing zinc and the phenolic preservative metacresol with phenoxyethanol as an antimicrobial agent and an amphiphilic acrylamide copolymer excipient for stability. A greater understanding is needed of the interplay between excipients, insulin association state, and stability in order to optimize this formulation. Here, we formulated insulin with different preservatives and stabilizing excipient concentrations using both insulin lispro and regular human insulin and assessed the insulin association states using analytical ultracentrifugation as well as formulation stability. We determined that phenoxyethanol is required to eliminate hexamers and promote a high monomer content even in a zinc-free lispro formulation. There is also a concentration dependent relationship between the concentration of polyacrylamide-based copolymer excipient and insulin stability, where a concentration greater than 0.1 g/mL copolymer is required for a mostly monomeric zinc-free lispro formulation to achieve stability exceeding that of Humalog in a stressed aging assay. Further, we determined that under the formulation conditions tested zinc-free regular human insulin remains primarily hexameric and is not at this time a promising candidate for rapid-acting formulations.

## INTRODUCTION

Insulin is an essential protein drug for over 20 million patients worldwide with type 1 diabetes.^1^ While insulin formulations have advanced tremendously in the last century, there is a continued push to develop ultra-rapid insulin formulations that would better mimic endogenous insulin secretion and enable improved automated insulin delivery in “artificial pancreas” devices.^2,3^ Yet, current rapid-acting analogues still fall short of the ultra-rapid response of insulin secreted from a healthy pancreas.^4–9^ Administering a formulation of insulin monomers would result in the fastest absorption rate from the subcutaneous space, but the insulin monomer is highly unstable in formulation,^10,11^ which has thus far prevented commercial use of this strategy. Understanding the role of formulation excipients on insulin association state and stability is an important step towards commercializing an ultra-fast insulin formulation.

Excipients are often overlooked as the “inactive” ingredients in pharmaceutical formulations. Yet, in many drug formulations excipients play a critical role in drug viability improving solubility, absorption, and stability, of the active ingredient.^12,13^ Insulin is naturally stored as a zinc-stabilized hexamer in healthy beta cells, and this has been replicated in most commercial insulin formulations to maintain insulin stability and shelf-life.^14^ Yet, while zinc-stabilized hexamers dissociate almost instantaneously when secreted into the blood, the hexamer dissociates much slower in the subcutaneous space due to lower dilution effects.^15^ As a result, exogenously delivered insulin exhibits delayed onset and a longer duration of action that is heavily dependent on the dissociation of the insulin hexamers first into dimers and then into the active monomers that are absorbed into the blood.

Efforts to reduce hexamer content by removing formulation zinc have been made in the rapid-acting formulation Apidra (Sanofi-Aventis), which is a zinc-free insulin analogue (glulisine); however, phenolic preservatives used as antimicrobial agents in all insulin formulations stabilize the R6 hexamer formation through hydrogen bonding.^16^ As a result Apidra pharmacokinetics/ pharmacodynamics are ultimately similar to those of zinc containing rapid-acting insulin formulations Humalog (Eli Lilly) and Novolog (Novo Nordisk).^17^

Recent work has identified a stable ultrafast insulin lispro formulation that isolates the insulin monomer by removing zinc and the phenolic preservative metacresol,^18,19^ and replacing them with phenoxyethanol as an antimicrobial agent and an amphiphilic acrylamide copolymer excipient for stability. This formulation contains about 70% insulin monomers, is stable twice as long as Humalog in a stressed aging test, demonstrates peak action in pigs of 10 minutes, and is predicted to have kinetics that are four times faster than Humalog in humans based on modelling.^19^ These results are extremely promising, but a greater understanding of the excipients is needed for formulation optimization before commercialization.

In this study, we aim to expand our understanding of preservative concentrations and insulin association state as well as optimize the concentration of polymeric excipient needed to achieve stability. Additionally, we look to explore the use of these excipients with regular human insulin to see if a similar formulation can be achieved as recombinant human insulin remains more affordable to both manufacture and obtain for patients. If excipients could be modulated to achieve an ultra-fast acting regular human insulin it would improve patient access to formulations.

## EXPERIMENTAL METHODS

### Materials

The amphiphilic acrylamide copolymer excipient poly(acryloylmorpholine77%-*co*-N-isopropylacrylamide_23%_) (MoNi) was prepared according to published protocols.^19^ Characterization of MoNi molecular weight can be found in Table S1. Humalog (Eli Lilly) and Humulin R (Eli Lilly) were purchased and used as received. For zinc-free lispro and zinc-free regular human insulin, Zinc(II) was removed from the commercial insulin formulations through competitive binding by addition of ethylenediaminetetraacetic acid (EDTA), which exhibits a dissociation binding constant approaching attomolar concentrations (K_D_~10^-18^ M).^20,21^ EDTA was added to formulations (4 molar equivalents with respect to zinc) to sequester zinc from the formulation and then lispro was isolated using PD MidiTrap G-10 gravity columns (GE Healthcare) to buffer exchange into water. The solution was then concentrated using Amicon Ultra 3K centrifugal units (Millipore) and reformulated with glycerol, phenoxyethanol, metacresol, methylparaben, and/or propylparaben in 10 mM phosphate buffer (pH=7.4) at an insulin concentration of 3.45 mg/mL (100 U/mL) (excipient concentrations specified in Table 1-2). For zinc-free formulations glycerol was added at 2.6 wt.% to be consistent with our previously reported monomeric lispro formulations,^18,19,22^ however a reduction of glycerol to 1.6% to match Humalog is not expected to affect insulin association states (Figure S1). All other reagents were purchased from Sigma-Aldrich unless otherwise specified.

**Table 1.** Excipient concentrations for insulin lispro formulations.

| **Formulation** | **Humalog** | **Lispro-1** | **Lispro-2** | **Lispro-3** | **Lispro-4** |
| --- | --- | --- | --- | --- | --- |
| ***Zinc*** | 0.0019 wt. % | - | - | - | - |
| ***Glycerol*** | 1.6 wt. % | 2.6 wt. % | 2.6 wt. % | 2.6 wt. % | 2.6 wt. % |
| ***Metacresol*** | 0.32 wt. % | 0.32 wt. % | 0.076 wt. % | - | - |
| ***Phenoxyethanol*** | - | - | 0.64 wt % | 0.85 wt. % | - |
| ***Methylparaben*** | - | - | - | - | 0.137 wt. % |
| ***Propylparaben*** | - | - | - | - | 0.015 wt. % |
| ***MoNi*** | - | 0.01 wt. % | 0.01 wt. % | 0.01 wt. % | 0.01 wt. % |

**Table 2.** Excipient concentrations for regular human insulin formulations.

| **Formulation** | **Humulin** | **RHI-1** | **RHI-2** | **RHI-3** | **RHI-4** |
| --- | --- | --- | --- | --- | --- |
| ***Zinc*** | 0.0019 wt. % | - | - | - | - |
| ***Glycerol*** | 1.6 wt. % | 2.6 wt. % | 2.6 wt. % | 2.6 wt. % | 2.6 wt. % |
| ***Metacresol*** | 0.32 wt. % | 0.32 wt. % | 0.076 wt. % | - | - |
| ***Phenoxyethanol*** | - | - | 0.64 wt % | 0.85 wt. % | - |
| ***Methylparaben*** | - | - | - | - | 0.137 wt. % |
| ***Propylparaben*** | - | - | - | - | 0.015 wt. % |
| ***MoNi*** | - | 0.01 wt. % | 0.01 wt. % | 0.01 wt. % | 0.01 wt. % |

### Analytical Ultracentrifugation

Insulin formulations were prepared at 3.45 mg/mL (100 U/mL) with excipients as described in Table 1-2. Analytical ultracentrifugation was performed by HTL Biosolutions. Briefly, samples were analyzed using a ProteomeLab XL-I analytical ultracentrifuge with 2-channel charcoal-epon centerpieces with 12 mm optical pathlength and an 8-hole An-50 Ti analytical rotor (20 °C, 45,000 rpm). Data were analyzed by HTL Biosolutions using SEDFIT version 16.2b and Gussi 1.3.1 software. SEDFIT (version 16.2b) developed by Peter Schuck at the N.I.H. was used for data analysis. The scans 1-200 were included for data analysis using the *c(s)* method. The *f/f_o_* values and meniscus position were fitted to find the best overall fit of the data for each sample. A maximum entropy regularization probability of 0.683 (1 σ) was used, and time-independent noise was removed. The continuous c(s) distribution model was used for data analysis of all samples as the signal from polymer is less than 3% of the total signal (the polymer concentration in each sample is 0.1 mg/mL and the insulin concentration is 3.5 mg/ mL).

The following parameter was used for the AUC-SV distribution analysis:

Partial specific volume: 0.73 mL/g

Density: 1.0000 mg/mL

Viscosity: 0.01002 P

Analytical ultracentrifugation is a useful method to determine insulin association states as sedimentation coefficients can be used to identify different association states and sedimentation velocity can be used to estimate the relative amounts of each species present. Since sedimentation coefficients are dependent on the molecular shape and mass of proteins, association state can be predicted when the monomer sedimentation coefficient is known. If we assume the aggregate shape is similar to that of the monomer then its stoichiometry, *N*, will be given by (*s_N_ /s_1_*)_3/2_, where *s_N_* is the sedimentation coefficient of the *N*-mer and *s_1_* is the sedimentation coefficient of monomer. Table S2 describes the general ranges of sedimentation coefficients provided by HTL Biosolutions (relative to the monomer sedimentation coefficient) used to assign association states. Observed sedimentation coefficients are dependent on solvent/buffer density, viscosity, and temperature, as well as transient species (those with lifetimes of less than ~1h) can cause a shift in sedimentation coefficients. When necessary, association state was assigned by rounding to the nearest range of sedimentation coefficients (assignment can be found in Table S3 and S4).

### In vitro stability

Aggregation assays used to evaluate stability were adapted from Webber et al.^23^ Briefly, formulations were aliquoted 150 μL per well (n = 3/group) in a clear 96-well plate and sealed with optically clear and thermally stable seal (VWR). The plate was incubated in a microplate reader (BioTek SynergyH1 microplate reader) at 37 °C with continuous agitation (567 cpm). Absorbance readings were taken every 10 minutes at 540 nm for the duration of the experiment. The formation of insulin aggregates leads to light scattering and a reduction in the transmittance of samples (time to aggregation = time to 10% change in transmittance).

### Statistics

All data is shown as mean ± s.e.m. unless specified otherwise. For time to aggregation data, a one-way ANOVA with a Tukey-Kramer correction for multiple comparisons was performed in GraphPad Prism 9, and each formulation was compared to all other formulations. Adjusted p values are listed in the supplemental information Table S5-8 (α < 0.05).

## RESULTS & DISCUSSION

### Excipients and insulin association state

Previous research using SEC-MALS has indicated that phenoxyethanol increases the proportion of monomers in formulation compared to metacresol.^18,19^ Unfortunately, measurements using SEC-MALS require samples to be formulated at very high concentrations (approximately 10-fold higher than formulation concentrations) so that the insulin is at relevant concentrations when it passes the detector. Further, shear forces as insulin travels along the column could result in dissociation of higher order species. Thus, we chose to further explore the effects of antimicrobial preservatives on insulin association state using analytical ultracentrifugation. Analytical ultracentrifugation is a valuable method for examining association states, because sedimentation coefficients can be used to identify different oligomers and the relative amounts of each species can be determined using sedimentation velocity (Table S3, S4).

As expected, insulin lispro formulations without zinc, regardless of preservative, had fewer hexamers compared to commercial Humalog (Figure 2). Metacresol as the only preservative in a zinc-free lispro formulation (Lispro-1) results in the lowers monomer content at 39%. The metacresol-phenoxyethanol combination (Lispro-2) and phenoxyethanol (Lispro-3) resulted in similar insulin lispro association states, both with 57% monomers. It was expected that replacing metacresol with phenoxyethanol would result in greater monomer content. Metacresol stabilizes the R6 hexamer assembly via hydrogen bonding to a hydrophobic pocket between monomers in the insulin hexamer configuration.^16,24,25^ In contrast, for phenoxyethanol we hypothesize that the ethoxyether functionality on the phenoxy moiety results in significant steric hindrance that prevents effective binding within the hydrophobic pocket of the insulin hexamer. In fact, the ethoxy functionality may reduce insulin stability, and promote the monomer form.^26^ The preservative combination of methylparaben and propylparaben (Lispro-4) was successful at increasing monomer and dimer content, but had a greater percentage of hexamers (19% hexamers) compared to formulations containing phenoxyethanol (Lispro 2 and 3: 0-0.5% hexamers). This observation is consistent with previous observations that insulin can also form a hexamer with methylparaben.^24^ Glycerol was used as a tonicity agent for all formulations since it is regularly used in commercial formulations and resulted in the highest monomer content when previously tested with SEC-MALS.

**Figure 1.**
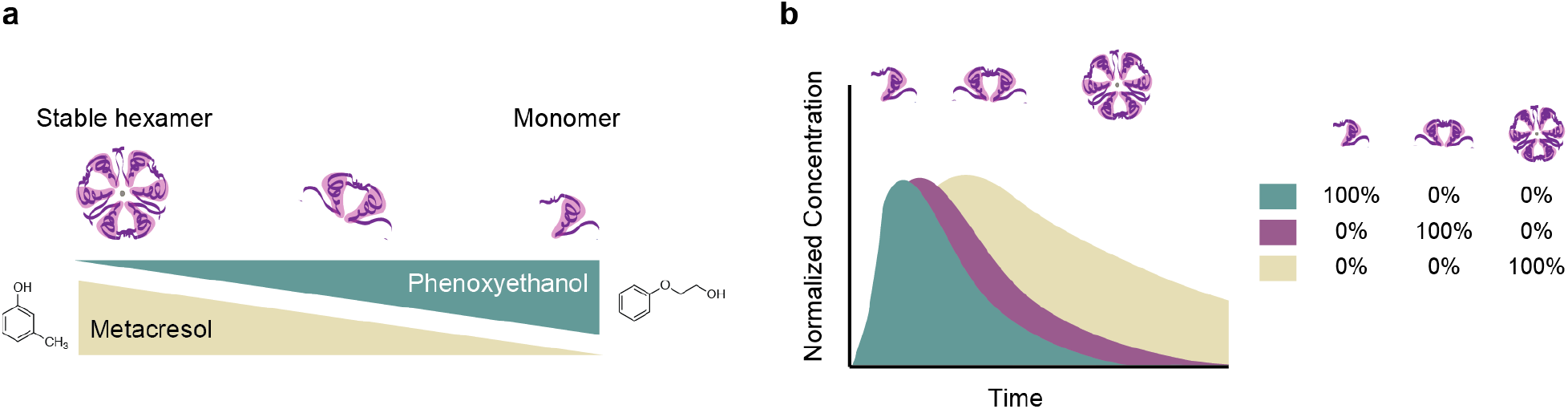
Schematic of insulin association states, preservatives and pharmacokinetics. A) Schematic showing how preservatives can promote different association states. The insulin hexamer is stable, while the insulin monomer is susceptible to rapid aggregation and amyloid fibril formation. Formulations containing high monomer counts require additional stabilizing agents. B) Schematic showing how insulin pharmacokinetics can be shifted by altering insulin association state. The insulin monomer is more rapidly absorbed into the blood. In contrast, subcutaneous administration of the insulin hexamer results in a delayed peak and longer duration of action.

**Figure 2.**
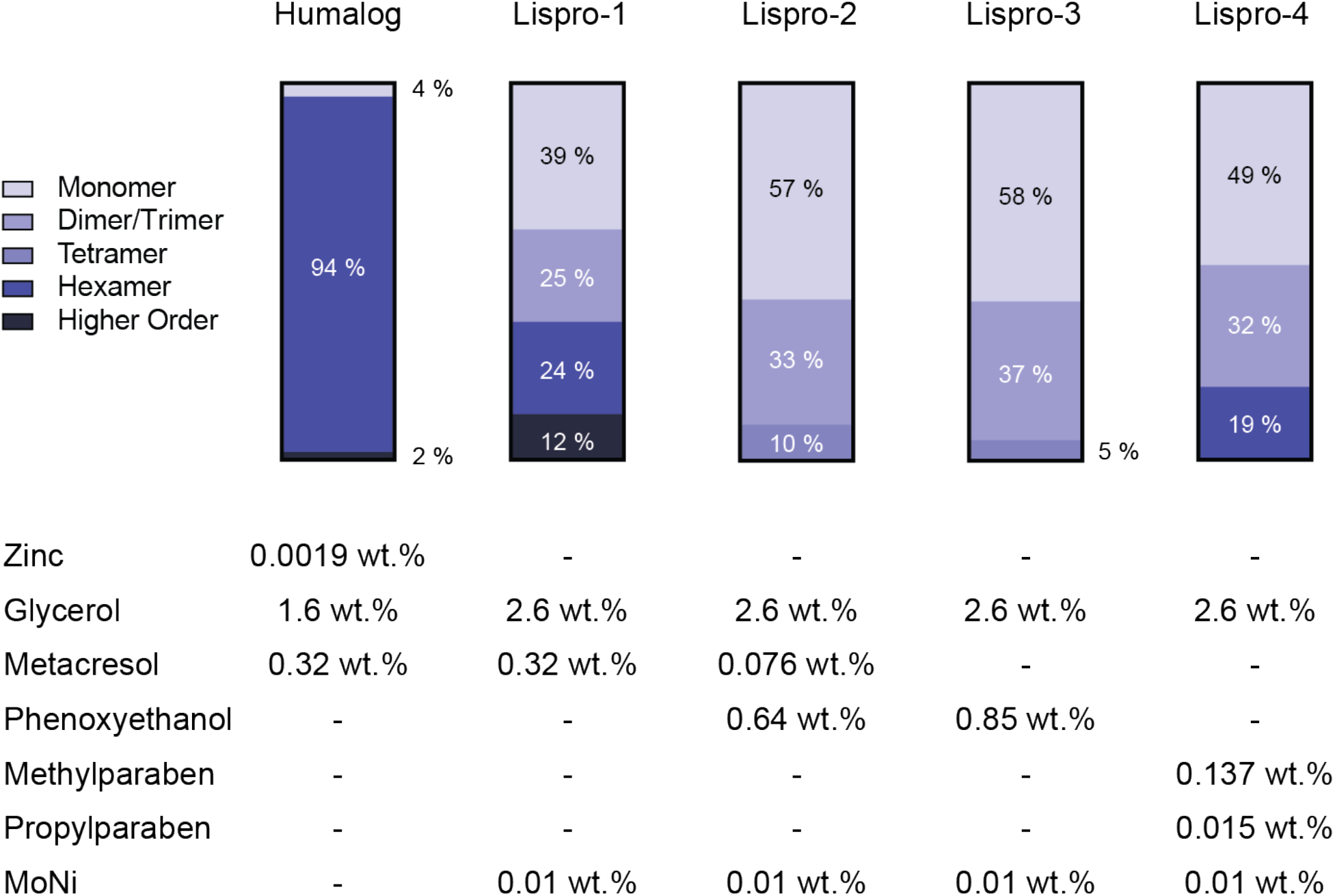
Insulin lispro association states. Using analytical ultracentrifugation, sedimentation coefficients were used to estimate the ratios of insulin association states when formulated with different antimicrobial preservatives in the absence of zinc. Commercial Humalog is shown as a control. It should be noted that higher ordered structures as reported here is thought to be reversible aggregates such as octamers, rather than high molecular weight polymer or amyloid fibril species.

In contrast to the lispro formulations, zinc-free regular human insulin resulted in low monomer content for all preservatives (Figure 3). The highest monomer content in zinc-free regular human insulin formulations was seen in the phenoxyethanol formulation (RHI-3), which contained 22% monomers, 8% dimers/trimers, and 70% hexamers. It stands out to us that there was not a higher dimer/trimer content observed in these zinc-free formulations. The primary difference in association state equilibria between rapid-acting insulin analogues and regular human insulin in the absence of zinc is the propensity for dimerization. Regular human insulin has 300-fold higher affinity for dimerization compared to insulin lispro.^27^ In contrast, the affinity for hexamer formation is only ~4-fold greater for regular human insulin.^27^ Future study may be needed to understand if other formulation components may promote the hexamer or higher ordered structures. It should be noted that higher ordered structures as reported here is thought to be reversible aggregates such as octamers, rather than high molecular weight polymer or amyloid fibril species. At present, the low monomer/dimer content observed in the tested formulations suggest that regular human insulin may not be well suited for the development of ultra-rapid insulin formulations.

**Figure 3.**
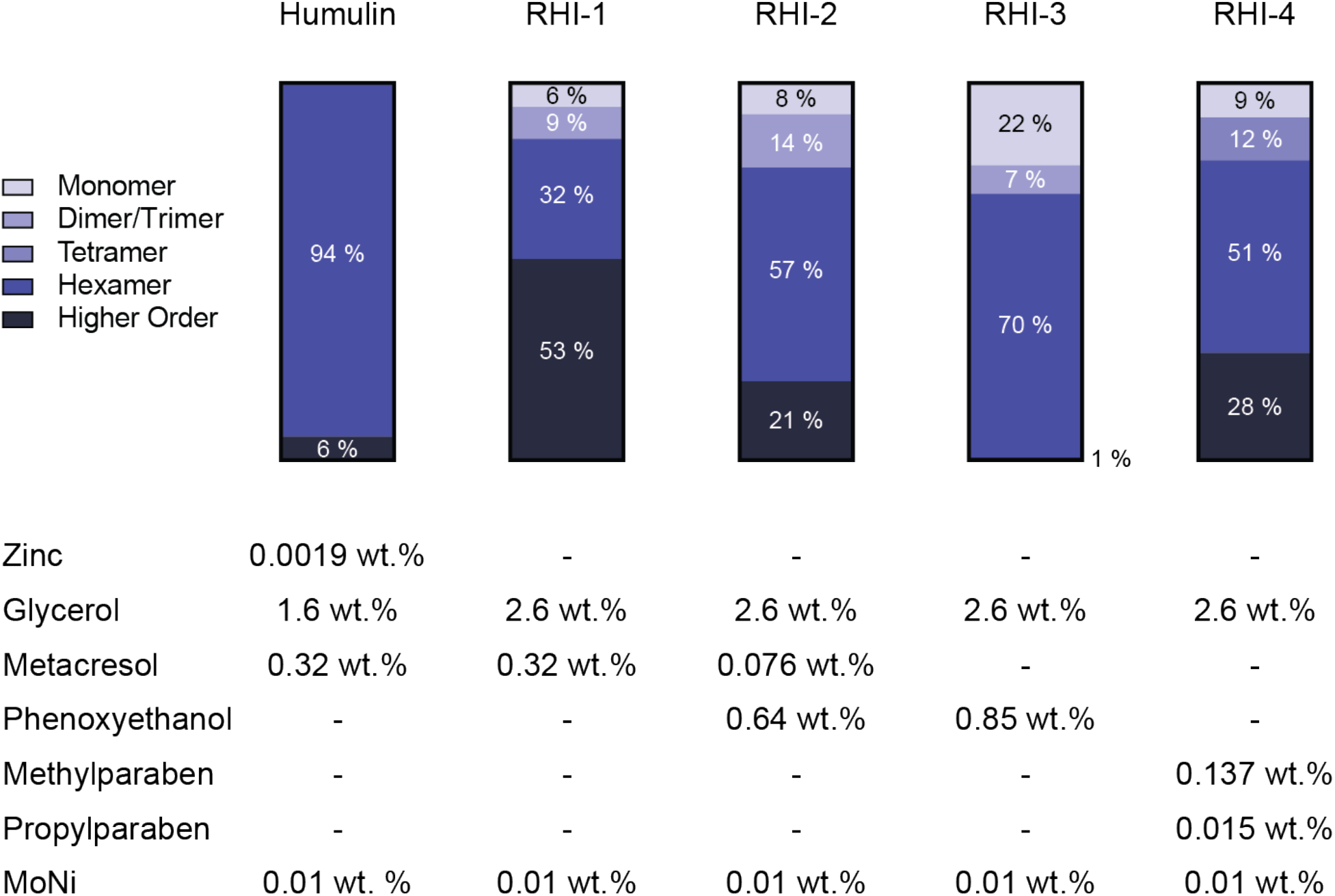
RHI association states. Using analytical ultracentrifugation, sedimentation coefficients were used to estimate the ratios of insulin association states when formulated with different antimicrobial preservatives in the absence of zinc. Commercial Humalog is shown as a control. It should be noted that higher ordered structures as reported here is thought to be reversible aggregates such as octamers, rather than high molecular weight polymer or amyloid fibril species.

### Excipients and formulation stability

The balance between the monomer content of an insulin formulation and stability remains a challenge in developing next-generation insulin formulations. Insulin aggregation typically occurs at hydrophobic interfaces, such as the air-water interface, where insulin-insulin interactions can nucleate aggregation. In this study, we use our previously reported copolymer excipient, poly(acryloylmorpholine77%-co-N-ispropylacrylamide23%) (MoNi), as a stabilizing agent in our zinc-free insulin formulations.^19,28^ As an amphiphilic polymer, we hypothesized that our MoNi excipient preferentially occupies the air-water interface where it can block or disrupt aggregation nucleating insulin-insulin interactions. This hypothesis is supported by our previous work where we show that the surface tension of insulin formulations containing the MoNi excipient are identical to formulations of the MoNi excipient alone in buffer, demonstrating that the MoNi species are dominating the interfacial interactions.^28^ Interfacial complex rheology measurements corroborate this observation, demonstrating that the addition of polymer disrupts insulin surface interactions and concomitant insulin-insulin aggregation events.^28^ Here, we look to evaluate the stability of our formulations using a stressed aging assay to measure insulin aggregation (Figure 4A-B). The formulations with the greatest monomer content (Lispro-3, RHI-3) were then tested with different concentrations of MoNi to assess what the optimal polymer concentration is required to attain sufficient stability for commercial viability (Figure 4C-D).

**Figure 4.**
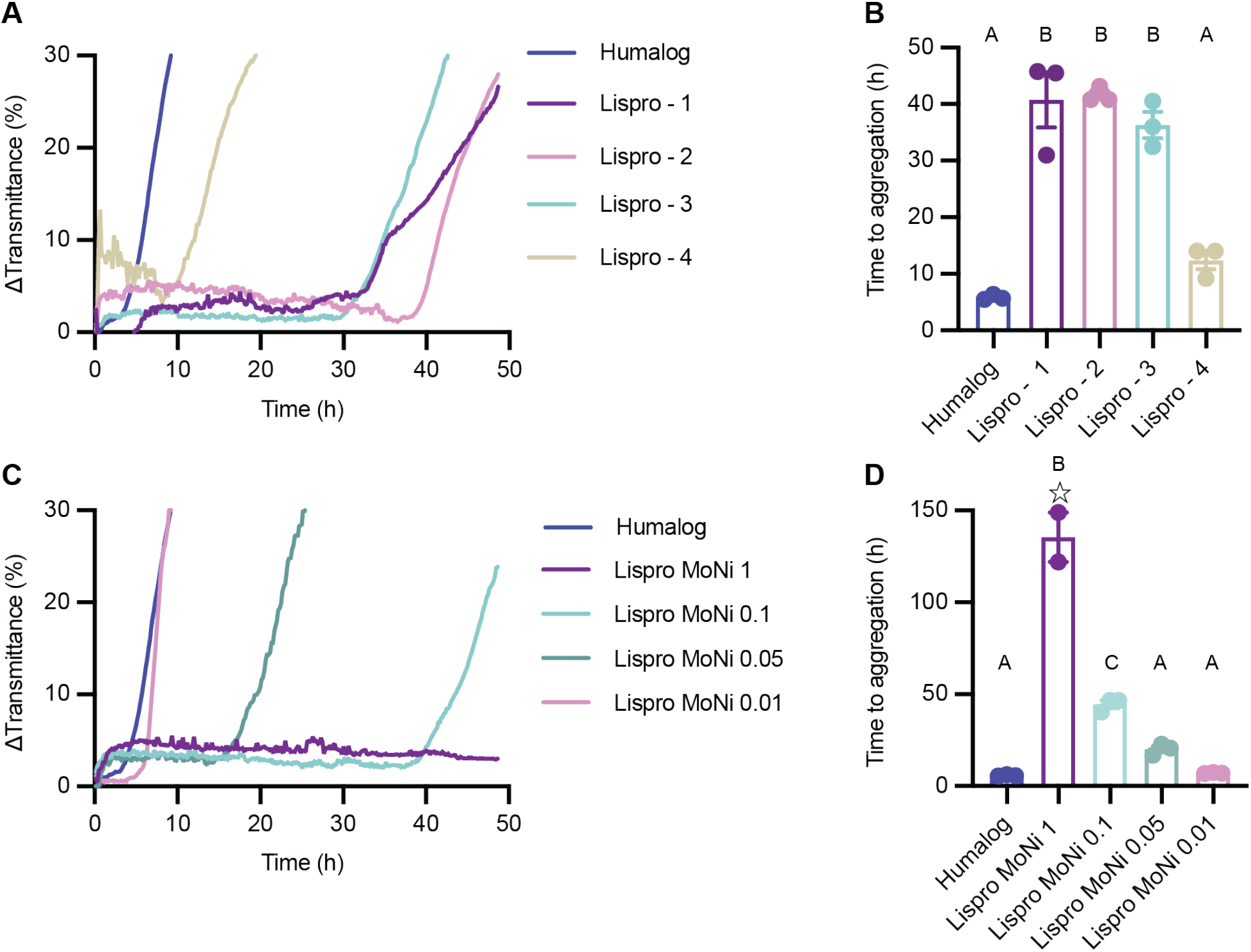
Insulin lispro formulation stability. A) Change in transmittance traces and B) Time to aggregation of formulations with different antimicrobial preservatives: Lispro - 1 (metacresol), Lispro - 2 (metacresol + phenoxyethanol), Lispro - 3 (phenoxyethanol), Lispro - 4 (methylparaben + propylparaben). C) Change in transmittance traces and D) Time to aggregation of phenoxyethanol formulation (Lispro - 3) with different concentrations of MoNi polymer excipient. Comparison of stability by aggregation times (t_A_), defined as the time to a change in transmittance (λ = 540 nm) of 10% or greater following stressed aging (i.e., continuous agitation at 37 °C). Data shown are average transmittance traces for n = 3 samples per group and error bars are standard error mean. Each ☆ represents a sample that did not aggregate before the assay ended at 150 hours. A one-way ANOVA with a Tukey-Kramer correction for multiple comparisons was performed in GraphPad Prism 9, and each formulation was compared to all other formulations. Formulations connected by the same letter label are not significantly different. Adjusted p values are listed in the supplemental information Table S5-6. α < 0.05.

Formulations containing our MoNi polymer excipient (0.1 mg/mL polymer) with metacresol and/ or phenoxyethanol (Lispro-1 t_A_=41 ± 8 h, P=0.0001; Lispro-2 t_A_=42 ± 2 h, P=0.0001; Lispro-3 t_A_=36 ± 4 h, P=0.0001) exhibited extended stability in our stressed aging assay compared to commercial Humalog (t_A_=5.8 ± 0.4 h) (Figure 4A-B). Similar times to aggregation were seen between commercial Humalog and the paraben containing formulation (Lispro-4 t_A_=12 ± 3 h, P=0.2666). Testing the stability of Lispro-3 with different concentrations of MoNi (Figure 4C-D), showed that the stability of formulations with high monomer content is dependent on polymer concentration. Formulations containing 1 mg/mL (140 ± 20 hours; P=0.0001) and 0.1 mg/mL (45 ± 4 hours, P=0.0001) MoNi demonstrated extended stability compared to Humalog. We also observed that the order of magnitude increase in MoNi concentration to 1 mg/mL from our standard MoNi concentration of 0.1 mg/mL resulted in almost 100 hours longer stability (P=0.0001). Decreasing the MoNi concentration to 0.05 mg/mL lowers the time to aggregation to 20 ± 3 hours, aggregating approximately twice as fast as the the formulation with 0.1 mg/mL polymer excipient (P=0.005). However, compared to commercial Humalog the time to aggregation for 0.05 mg/mL (T_A_: 20 ± 3 hours; p=0.061), 0.01 mg/mL (T_A_: 7.1 ± 0.3 hours; p=0.999), and 0 mg/mL (T_A_: 9 ± 2 hours; p=0.965) are not statistically different, indicating that lower concentrations of MoNi provide limited stability advantages. Since insulin aggregation is a stochastic process, it becomes difficult to differentiate between formulations that have very rapid times to aggregation. It is expected that no polymer in a mostly monomeric insulin formulation like Lispro-3 should be much less stable than commercial Humalog. While outside the scope of the present study, future studies will aim to explore covalent dimer formation and high molecular weight polymer formation in formulations to better differentiate the stability of formulations with lower polymer content compared to commercial Humalog.

Zinc-free formulations that were prepared with regular human insulin were more stable than zinc-free lispro formulations (Figure 5). Lispro-2, the most stable zinc-free lispro formulation, aggregated after 42 ± 2 h, while RHI-3, the most stable zinc-free RHI formulation, aggregated in 140 ± 10 h. This is likely due to reduced monomer content and the greater percentage of stable insulin hexamers in formulation observed in the analytical ultracentrifugation data. All four zinc-free formulations with MoNi polymer excipient (0.1 mg/mL polymer) demonstrated extended stability compared to commercial Humalog (RHI-1 t_A_=90 ± 20 h, P=0.0005; RHI-2 t_A_=140 ± 13 h, P=0.0001; RHI-3 t_A_=140 ± 10 h, P=0.0001; RHI-4 t_A_=100 ± 20 h, P=0.0001; Humalog t_A_=5.8 ± 0.4 h). No statistical difference was observed between the zinc-free regular human insulin formulations. When different concentrations of MoNi were tested in combination with RHI-3, all formulations also had greater stability compared to Humalog.

**Figure 5.**
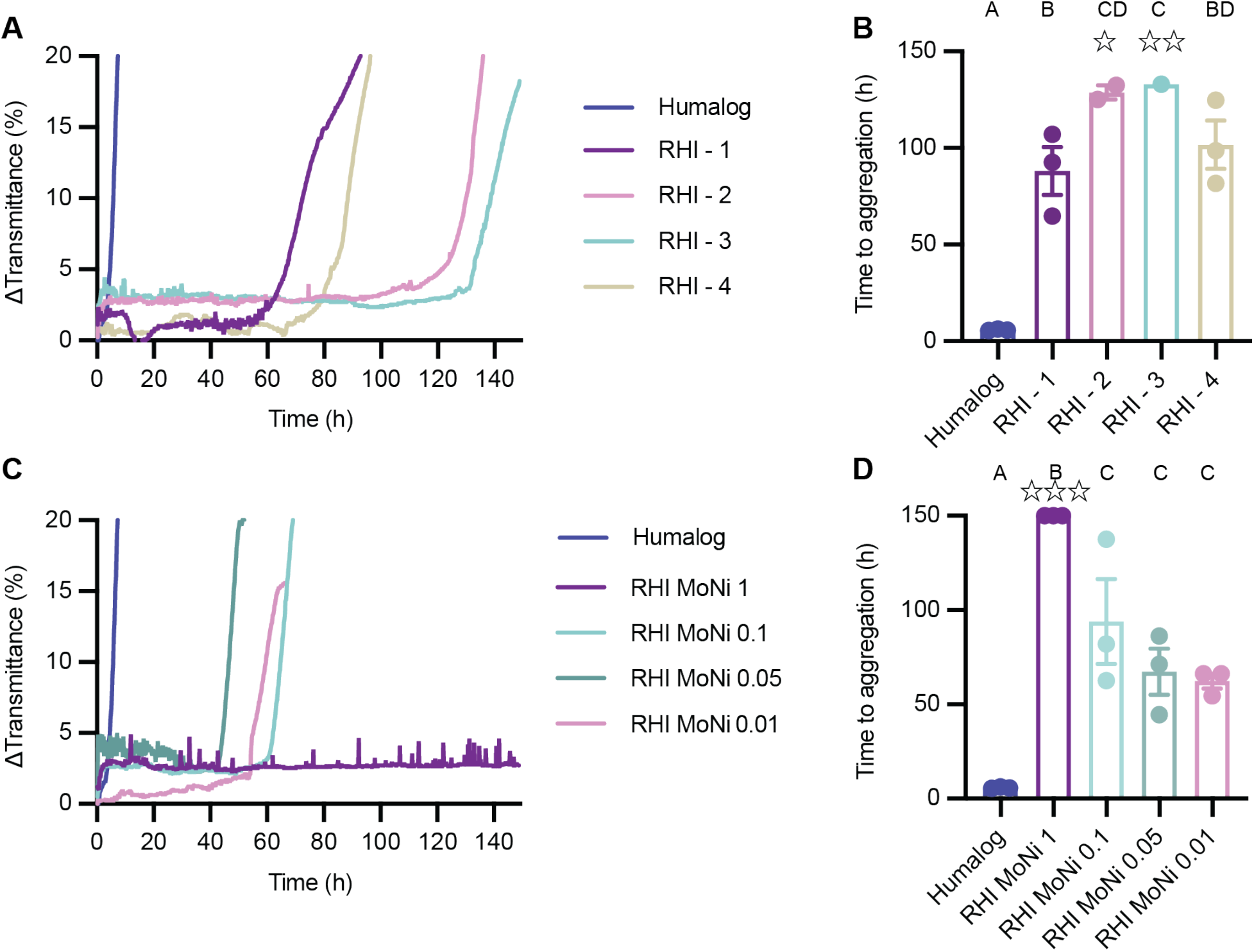
RHI formulation stability. A) Change in transmittance traces and B) Time to aggregation of formulations with different antimicrobial preservatives: RHI - 1 (metacresol), RHI - 2 (metacresol + phenoxyethanol), RHI - 3 (phenoxyethanol), RHI - 4 (methylparaben + propylparaben). C) Change in transmittance traces and D) Time to aggregation of phenoxyethanol formulation (RHI - 3) with different concentrations of MoNi polymer excipient. Comparison of stability by aggregation times (tA), defined as the time to a change in transmittance (λ = 540 nm) of 10% or greater following stressed aging (i.e., continuous agitation at 37 °C). Data shown are average transmittance traces for n = 3 samples per group and error bars are standard error mean. Each ☆ represents a sample that did not aggregate before the assay ended at 150 hours. A one-way ANOVA with a Tukey-Kramer correction for multiple comparisons was performed in GraphPad Prism 9, and each formulation was compared to all other formulations. Formulations connected by the same letter label are not significantly different. Adjusted p values are listed in the supplemental information Table S7-8. α < 0.05.

## CONCLUSION

Evaluation of the association state and stability of different zinc-free insulin lispro and regular human insulin formulations revealed that the use of phenoxyethanol as an antimicrobial excipient remains a key component to achieving high monomer content formulations with elimination of the presence of insulin hexamers — critical characteristics of a ultrafast insulin formulation. Unfortunately these studies suggest that under the tested formulation conditions, zinc-free regular human insulin remains primarily hexameric and is not suitable for developing a ultrafast insulin formulation, highlighting that insulin analogs such as Lispro and Aspart must be used. Stressed aging assays identify that the stability enabled by amphiphilic copolymer excipient MoNi is concentration dependent and concentrations of 0.1 mg/mL are required to extend the stability of a mostly monomeric zinc-free lispro formulation compared to Humalog. Overall, these results give us insight on how these excipients modify formulation properties and in the future they could be used to tune insulin pharmacokinetics. Future study developing models to directly predict insulin pharmacokinetics from insulin association state could be powerful for optimizing formulations.

## Acknowledgments

This work was funded in part by NIDDK R01 (NIH grant #R01DK119254), and a Pilot and Feasibility funding from the Stanford Diabetes Research Center (NIH grant #P30DK116074) and the American Diabetes Association Grant (1-18-JDF-011). Support is also provided by the Stanford Maternal and Child Health Research Institute through the SPARK Translational Research Program. C.L.M. was supported by the NSERC Postgraduate Scholarship and the Stanford BioX Bowes Graduate Student Fellowship. J.L.M was supported Department of Defense NDSEG Fellowship and by a Stanford Graduate Fellowship. The authors thank the Stanford Animal Diagnostic Lab and the Veterinary Service Centre staff for their technical assistance.

## Author Contributions

C.L.M. and E.A.A designed the experiments. C.L.M. and L.T.N. performed the experiments. J.L.M. synthesized the polymers. C.L.M. analyzed the data. C.L.M. and E.A.A. wrote the manuscript. All authors revised the manuscript.

## Data Availability

All data supporting the results in this study are available within the Article and its Supplementary Information. Raw data files are available from the corresponding author upon reasonable request.

## References

1. International Diabetes Federation IDF Diabetes Atlas, 9th edn.; https://www.diabetesatlas.org., 2019.

2. Cengiz, E., Undeniable need for ultrafast-acting insulin: the pediatric perspective. J. Diabetes Sci. Technol. 2012, 6, 797–801.

3. Gingras, V.; Taleb, N.; Roy-Fleming, A.; Legault, L.; Rabasa-Lhoret, R., The challenges of achieving postprandial glucose control using closed-loop systems in patients with type 1 diabetes. Diabetes Obes. Metab. 2018, 20, 245–256.

4. Polonsky, K. S.; Given, B. D.; Van Cauter, E., Twenty-four-hour profiles and pulsatile patterns of insulin secretion in normal and obese subjects. J. Clin. Invest. 1988, 81, 442–448.

5. Tillil, H.; Shapiro, E. T.; Miller, M. A.; Karrison, T.; Frank, B. H.; Galloway, J. A.; Rubenstein, A. H.; Polonsky, K. S., Dose-dependent effects of oral and intravenous glucose on insulin secretion and clearance in normal humans. Am. J. Physiol. 1988, 254, E349–57.

6. Heptulla, R. A.; Rodriguez, L. M.; Bomgaars, L.; Haymond, M. W., The Role of Amylin and Glucagon in the Dampening of Glycemic Excursions in Children With Type 1 Diabetes. Diabetes 2005, 54, 1100.

7. Saad, A.; Dalla Man, C.; Nandy, D. K.; Levine, J. A.; Bharucha, A. E.; Rizza, R. A.; Basu, R.; Carter, R. E.; Cobelli, C.; Kudva, Y. C.; Basu, A., Diurnal Pattern to Insulin Secretion and Insulin Action in Healthy Individuals. Diabetes 2012, 61, 2691.

8. Heise, T.; Zijlstra, E.; Nosek, L.; Rikte, T.; Haahr, H., Pharmacological properties of faster-acting insulin aspart vs insulin aspart in patients with type 1 diabetes receiving continuous subcutaneous insulin infusion: A randomized, double-blind, crossover trial. Diabetes Obes. Metab. 2017, 19, 208–215.

9. Senior, P.; Hramiak, I., Fast-Acting Insulin Aspart and the Need for New Mealtime Insulin Analogues in Adults With Type 1 and Type 2 Diabetes: A Canadian Perspective. Can. J. Diabetes 2019, 43, 515–523.

10. Hua, Q. X.; Weiss, M. A., Mechanism of insulin fibrillation: the structure of insulin under amyloidogenic conditions resembles a protein-folding intermediate. J. Biol. Chem. 2004, 279, 21449–60

11. Woods, R. J.; Alarcon J Fau - McVey, E.; McVey E Fau - Pettis, R. J.; Pettis, R. J., Intrinsic fibrillation of fast-acting insulin analogs. J. Diabetes Sci. Technol. 2012, *6*. 265–276.

12. Kamerzell, T. J.; Esfandiary, R.; Joshi, S. B.; Middaugh, C. R.; Volkin, D. B., Protein-excipient interactions: mechanisms and biophysical characterization applied to protein formulation development. Adv. Drug Deliv. Rev. 2011, 63, 1118–1159.

13. Rayaprolu, B. M.; Strawser, J. J.; Anyarambhatla, G., Excipients in parenteral formulations: selection considerations and effective utilization with small molecules and biologics. Drug Dev. Ind. Pharm. 2018, 44, 1565–1571.

14. Xu, Y; Yan Y. Fau - Seeman, D.; Seeman D Fau - Sun, L.; Sun L Fau - Dubin, P. L.; Dubin, P. L., Multimerization and aggregation of native-state insulin: effect of zinc. Langmuir. 2012, 28, 579–586

15. Gast, K.; Schüler, A.; Wolff, M.; Thalhammer, A.; Berchtold, H.; Nagel, N.; Lenherr, G.; Hauck, G.; Seckler, R., Rapid-Acting and Human Insulins: Hexamer Dissociation Kinetics upon Dilution of the Pharmaceutical Formulation. Pharm. Res. 2017, 34, 2270–2286.

16. Teska, B. M.; Alarcón, J.; Pettis, R. J.; Randolph, T. W.; Carpenter, J. F., Effects of phenol and meta-cresol depletion on insulin analog stability at physiological temperature. J. Pharm. Sci. 2014, 103, 2255–2267.

17. Home, P. D., The pharmacokinetics and pharmacodynamics of rapid-acting insulin analogues and their clinical consequences. Diabetes Obes. Metab. 2012, 14, 780–788.

18. Maikawa, C. L.; Smith, A. A. A.; Zou, L.; Meis, C. M.; Mann, J. L.; Webber, M. J.; Appel, E. A., Stable Monomeric Insulin Formulations Enabled by Supramolecular PEGylation of Insulin Analogues. Adv. Ther. 2019, 75, 1900094.

19. Mann, J. L.; Maikawa, C. L.; Smith, A. A. A.; Grosskopf, A. K.; Baker, S. W.; Roth, G. A.; Meis, C. M.; Gale, E. C.; Liong, C. S.; Correa, S.; Chan, D.; Stapleton, L. M.; Yu, A. C.; Muir, B.; Howard, S.; Postma, A.; Appel, E. A., An Ultra-fast Insulin Formulation Enabled by High Throughput Screening of Polymeric Excipients. Sci. Transl. Med. 2020, 12, eaba6676.

20. Berthon, G., Handbook of metal-ligand interactions in biological fluids. Bioinorganic chemistry. Marcel Dekker: New York, 1995.

21. Waters, R. S.; Bryden, N. A.; Patterson, K. Y.; Veillon, C.; Anderson, R. A., EDTA chelation effects on urinary losses of cadmium, calcium, chromium, cobalt, copper, lead, magnesium, and zinc. Biol. Trace Elem. Res. 2001, 83, 207–21.

22. Maikawa, C. L.; Chen, P. C.; Vuong, E. T.; Nguyen, L. T.; Mann, J. L.; d’Aquino, A. I.; Lal, R. A.; Maahs, D. M.; Buckingham, B. A.; Appel, E. A. Ultra-Fast Insulin–Pramlintide Co-Formulation for Improved Glucose Management in Diabetic Rats. Adv. Sci. 2021, 8, 2101575.

23. Webber, M. J.; Appel, E. A.; Vinciguerra, B.; Cortinas, A. B.; Thapa, L. S.; Jhunjhunwala, S.; Isaacs, L.; Langer, R.; Anderson, D. G., Supramolecular PEGylation of biopharmaceuticals. Proc. Natl. Acad. Sci. U. S. A. 2016, 113, 14189–14194.

24. Whittingham, J. L.; Chaudhuri, S.; Dodson, E. J.; Moody, P. C. E.; Dodson, G. G. X-ray Crystallographic Studies on Hexameric Insulins in the Presence of Helix-Stabilizing Agents, Thiocyanate, Methylparaben and Phenol. Biochem. 1995, 34, 15553–15563.

25. Whittingham, J. L.; Edwards, D. J.; Antson, A. A.; Clarkson, J. M.; Dodson, G. G. Interactions of Phenol and m-Cresol in the Insulin Hexamer, and Their Effect on the Association Properties of B28 Pro → Asp Insulin Analogues. Biochem. 1998, 37, 11516–11523.

26. Banerjee, P.; Mondal, S.; Bagchi, B. Effect of ethanol on insulin dimer dissociation. J. Chem. Phys. 2019, 150, 084902.

27. Brems, D. N.; Alter, L. A.; Beckage, M. J.; Chance, R. E.; DiMarchi, R. D.; Green, L. K.; Long, H. B.; Pekar, A. H.; Shields, J. E.; Frank, B. H., Altering the association properties of insulin by amino acid replacement. Protein Eng. Des. Sel. 1992, 5, 527–533.

28. Maikawa, C. L.; Mann, J. L.; Kannan, A.; Meis, C. M.; Grosskopf, A. K.; Ou, B. S.; Autzen, A. A. A.; Fuller, G. G.; Maahs, D. M.; Appel, E. A. Engineering Insulin Cold Chain Resilience to Improve Global Access. Biomacromolecules. 2021, 22, 3386–3395.

