## Supplemental Information for "Formulation excipients and their role in insulin stability and association state in formulation"

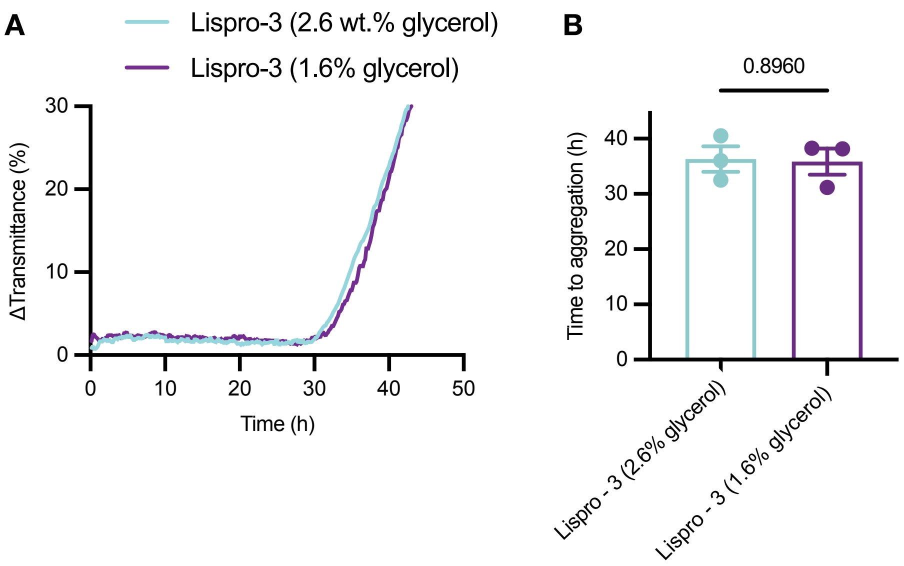


**Figure S1. Stability of Lispro-3 with different glycerol concentrations.** Here the Lispro-3 formulation only differs by the amount of glycerol added. A change in glycerol concentration from 1.6 wt.% as is common in commercial formulations to 2.6 wt.% does not have an effect on formulation stability. A) Change in transmittance traces and B) Time to aggregation. Time to aggregation was compared using a two-tailed t-test ($\alpha<0.05$) in GraphPad Prism 9.

**Table S1. MoNi copolymer excipient characterization.**

| Carrier Monomer | wt.% (Target) | wt.% by NMR (Exp) | Dopant Monomer | wt.% (Target) | wt.% by NMR (Exp) | $M_n$[a] (Da) | $M_w$[a] (Da) | Đ[a] |
|---|---|---|---|---|---|---|---|---|
| Acryloylmorpholine (Mo) | 77 | 74.5 [b] | N-isopropylacrylamide (Ni) | 23 | 25.5 [b] | 3200 | 3800 | 1.19 |

[a] Determined using Size Exclusion Chromatography calibrated using polyethylene glycol samples.

[b] Weight percentages difficult to determine due to overlapping spectra. Weight percentages estimated from post-precipitated NMR spectra by measuring the more resolved left half of the peak of N-isopropylacrylamide (δ= 4.0, 0.5 H), doubling it, and subtracting it from the unresolved peaks of Mo and Ni (δ= 3.2-4.2, 7H (Mo) 1H (Ni)).

**Table S2. Sedimentation Coefficients Corresponding to Insulin Association States**

| **Insulin Association State** | **Ratio of oligomer sedimentation coefficient to monomer sedimentation coefficient*** |
|---|---|
| Monomer | ~1 |
| Dimer/Trimer | 1.45-1.86 |
| Tetramer | 2-2.26 |
| Hexamer | 2.67-2.97 |
| Higher Order (Octamer+) | > 3.46 |

* These were provided by HTL Biosolutions as “rule of thumb” values. Assignment of association states were rounded to the nearest range of sedimentation coefficients

**Table S3. Ratio of Insulin Lispro Association States by Insulin Sedimentation Coefficients**

| | | | | | |
|---|---|---|---|---|---|
| | | | **Humalog** | | |
| **sed c(s)** | 1.14 | 2.86 | 5.83 | 8.23 | 8.94 |
| **% peak** | 3.6 | 94.3 | 1.1 | 0.4 | 0.6 |
| **classification** | Monomer | Hexamer | Higher Order | Higher Order | Higher Order |
| | | | **Lispro-1** | | |
| **sed c(s)** | 0.85 | 1.84 | 3.03 | 3.58 | >3.6 |
| **% peak** | 38.9 | 24.5 | 24.5 | 12 | 0.1 |
| **classification** | Monomer | Dimer/Trimer | Hexamer | Higher Order | Higher Order |
| | | | **Lispro-2** | | |
| **sed c(s)** | 0.89 | 1.62 | 2.37 | | |
| **% peak** | 57.3 | 33.1 | 9.6 | | |
| **classification** | Monomer | Dimer/Trimer | Tetramer | | |
| | | | **Lispro-3** | | |
| **sed c(s)** | 0.81 | 1.4 | 2.3 | >2.3 | |
| **% peak** | 57.8 | 36.8 | 4.9 | 0.5 | |
| **classification** | Monomer | Dimer/Trimer | Tetramer | Hexamer | |
| | | | **Lispro-4** | | |
| **sed c(s)** | 0.91 | 1.57 | 2.57 | >2.6 | |
| **% peak** | 48.4 | 32.3 | 19.3 | 0.1 | |
| **classification** | Monomer | Dimer/Trimer | Hexamer | Hexamer | |

**Table S4. Ratio of Regular Human Insulin Association States by Insulin Sedimentation Coefficients**

| | | | | | | |
|---|---|---|---|---|---|---|
| | | | **Humulin** | | | |
| **sed c(s)** | 2.8 | 5.78 | 9.14 | >9.2 | | |
| **% peak** | 93.81 | 3.29 | 0.9 | 2 | | |
| **classification** | Hexamer | Higher Order | Higher Order | Higher Order | | |
| | | | **RHI-1** | | | |
| **sed c(s)** | 0.83 | 1.85 | 3.1 | 3.61 | 4.37 | |
| **% peak** | 6.39 | 8.44 | 31.9 | 11.36 | 41.93 | |
| **classification** | Monomer | Dimer/Trimer | Hexamer | Higher Order | Higher Order | |
| | | | **RHI-2** | | | |
| **sed c(s)** | 1.02 | 1.86 | 2.83 | 3.37 | | |
| **% peak** | 8.36 | 14.09 | 56.7 | 20.86 | | |
| **classification** | Monomer | Dimer/Trimer | Hexamer | Higher Order | | |
| | | | **RHI-3** | | | |
| **sed c(s)** | 0.77 | 1.65 | 2.69 | >2.7 | | |
| **% peak** | 21.9 | 7.48 | 69.9 | 0.7 | | |
| **classification** | Monomer | Dimer/Trimer | Hexamer | Hexamer | | |
| | | | **RHI-4** | | | |
| **sed c(s)** | 0.87 | 1.96 | 2.85 | 3.06 | 3.71 | >3.8 |
| **% peak** | 9.17 | 11.38 | 26.89 | 24.07 | 28.36 | 0.1 |
| **classification** | Monomer | Tetramer | Hexamer | Hexamer | Higher Order | Higher Order |

**Table S5. Adjusted P Values Figure 4B**

| Comparison | Mean difference | Adjusted P Value |
|---|---|---|
| Lispro - 1 vs. Lispro - 2 | -0.8200 | 0.9993 |
| Lispro - 1 vs. Lispro - 3 | 4.433 | 0.7355 |
| Lispro - 1 vs. Humalog | 34.94 | <0.0001 |
| Lispro - 1 vs. Lispro - 4 | 28.37 | 0.0001 |
| Lispro - 2 vs. Lispro - 3 | 5.253 | 0.6083 |
| Lispro - 2 vs. Humalog | 35.76 | <0.0001 |
| Lispro - 2 vs. Lispro - 4 | 29.19 | <0.0001 |
| Lispro - 3 vs. Humalog | 30.50 | <0.0001 |
| Lispro - 3 vs. Lispro - 4 | 23.93 | 0.0004 |
| Humalog vs. Lispro - 4 | -6.570 | 0.4124 |

**Table S6. Adjusted P values for Figure 4D**

| Comparison | Mean difference | Adjusted P Value |
|---|---|---|
| Lispro MoNi 1 vs. Lispro MoNi 0.1 | 91.00 | <0.0001 |
| Lispro MoNi 1 vs. Lispro MoNi 0.05 | 115.4 | <0.0001 |
| Lispro MoNi 1 vs. Lispro MoNi 0.01 | 128.4 | <0.0001 |
| Lispro MoNi 1 vs. Lispro MoNi 0 | 126.8 | <0.0001 |
| Lispro MoNi 1 vs. Humalog | 129.7 | <0.0001 |
| Lispro MoNi 0.1 vs. Lispro MoNi 0.05 | 24.43 | 0.0050 |
| Lispro MoNi 0.1 vs. Lispro MoNi 0.01 | 37.45 | 0.0001 |
| Lispro MoNi 0.1 vs. Lispro MoNi 0 | 35.78 | 0.0002 |
| Lispro MoNi 0.1 vs. Humalog | 38.67 | 0.0001 |
| Lispro MoNi 0.05 vs. Lispro MoNi 0.01 | 13.01 | 0.1792 |
| Lispro MoNi 0.05 vs. Lispro MoNi 0 | 11.35 | 0.2867 |
| Lispro MoNi 0.05 vs. Humalog | 14.24 | 0.1242 |
| Lispro MoNi 0.01 vs. Lispro MoNi 0 | -1.667 | 0.9993 |
| Lispro MoNi 0.01 vs. Humalog | 1.223 | 0.9998 |
| Lispro MoNi 0 vs. Humalog | 2.890 | 0.9908 |

**Table S7. Adjusted P values for Figure 5B**

| Comparison | Mean difference | Adjusted P Value |
|---|---|---|
| Humalog vs. RHI - 1 | -82.27 | 0.0005 |
| Humalog vs. RHI - 2 | -130.0 | <0.0001 |
| Humalog vs. RHI - 3 | -138.5 | <0.0001 |
| Humalog vs. RHI - 4 | -95.82 | 0.0001 |
| RHI - 1 vs. RHI - 2 | -47.73 | 0.0233 |
| RHI - 1 vs. RHI - 3 | -56.23 | 0.0084 |
| RHI - 1 vs. RHI - 4 | -13.55 | 0.8154 |
| RHI - 2 vs. RHI - 3 | -8.500 | 0.9578 |
| RHI - 2 vs. RHI - 4 | 34.18 | 0.1220 |
| RHI - 3 vs. RHI - 4 | 42.68 | 0.0433 |

**Table S8. Adjusted P values for Figure 5D**

| Comparison | Mean difference | Adjusted P Value |
|---|---|---|
| Humalog vs. RHI MoNi 1 | -144.2 | <0.0001 |
| Humalog vs. RHI MoNi 0.1 | -88.17 | 0.0008 |
| Humalog vs. RHI MoNi 0.05 | -61.55 | 0.0139 |
| Humalog vs. RHI MoNi 0.01 | -56.54 | 0.0244 |
| Humalog vs. RHI MoNi 0 | -144.2 | <0.0001 |
| RHI MoNi 1 vs. RHI MoNi 0.1 | 56.00 | 0.0260 |
| RHI MoNi 1 vs. RHI MoNi 0.05 | 82.62 | 0.0014 |
| RHI MoNi 1 vs. RHI MoNi 0.01 | 87.63 | 0.0008 |
| RHI MoNi 1 vs. RHI MoNi 0 | 0.000 | >0.9999 |
| RHI MoNi 0.1 vs. RHI MoNi 0.05 | 26.62 | 0.5102 |
| RHI MoNi 0.1 vs. RHI MoNi 0.01 | 31.63 | 0.3405 |
| RHI MoNi 0.1 vs. RHI MoNi 0 | -56.00 | 0.0260 |
| RHI MoNi 0.05 vs. RHI MoNi 0.01 | 5.010 | 0.9993 |
| RHI MoNi 0.05 vs. RHI MoNi 0 | -82.62 | 0.0014 |
| RHI MoNi 0.01 vs. RHI MoNi 0 | -87.63 | 0.0008 |
